# Complete substitution with modified nucleotides suppresses the early interferon response and increases the potency of self-amplifying RNA

**DOI:** 10.1101/2023.09.15.557994

**Authors:** Joshua E. McGee, Jack R. Kirsch, Devin Kenney, Elizabeth Chavez, Ting-Yu Shih, Florian Douam, Wilson W. Wong, Mark W. Grinstaff

## Abstract

Self-amplifying RNA (saRNA) will revolutionize vaccines and *in situ* therapeutics by enabling protein expression for longer duration at lower doses. However, a major barrier to saRNA efficacy is the potent early interferon response triggered upon cellular entry, resulting in saRNA degradation and translational inhibition. Substitution of mRNA with modified nucleotides (modNTPs), such as N1-methylpseudouridine (N1mΨ), reduce the interferon response and enhance expression levels. Multiple attempts to use modNTPs in saRNA have been unsuccessful, leading to the conclusion that modNTPs are incompatible with saRNA, thus hindering further development. Here, contrary to the common dogma in the field, we identify multiple modNTPs that when incorporated into saRNA at 100% substitution confer immune evasion and enhance expression potency. Transfection efficiency enhances by roughly an order of magnitude in difficult to transfect cell types compared to unmodified saRNA, and interferon production reduces by >8 fold compared to unmodified saRNA in human peripheral blood mononuclear cells (PBMCs). Furthermore, we demonstrate expression of viral antigens *in vitro* and observe significant protection against lethal challenge with a mouse-adapted SARS-CoV-2 strain *in vivo*. A modified saRNA vaccine, at 100-fold lower dose than a modified mRNA vaccine, results in a statistically improved performance to unmodified saRNA and statistically equivalent performance to modified mRNA. This discovery considerably broadens the potential scope of self-amplifying RNA, enabling entry into previously impossible cell types, as well as the potential to apply saRNA technology to non-vaccine modalities such as cell therapy and protein replacement.

## INTRODUCTION

The original discovery of the application of modified nucleotides (modNTPs) to mRNA by Karikó and Weissman revolutionized RNA medicine [1, 2]. This breakthrough enabled rapid development and deployment of mRNA vaccines against SARS-CoV-2, saving millions of lives. Chemically modified nucleotides enhance mRNA stability, transfection capability, and decrease immunogenicity [2–4]. Without modNTPs, detection of exogenous RNA activates toll-like receptors (TLRs) and triggers the production of type I interferons, resulting in translational shutoff and systemic inflammation. In the cytosol, retinoic acid-inducible gene 1 (RIG1) and RNA-dependent protein kinase (PKR) recognize RNA and trigger additional interferon production [4]. Clinically approved mRNA vaccines utilize N1-methylpseudouridine (N1mΨ) to mitigate these responses and improve efficacy. Although effective, the inherent short half-life of mRNA necessitates a large dose to be effective, which increases risk of adverse side effects and limits accessibility. Therefore, efforts to further enhance the expression and durability of RNA-based medicine at lower doses will unlock new therapeutic applications, improve tolerability, and expand global access.

Self-amplifying RNAs (saRNAs) undergo replication and amplification, inside the cell, to afford robust and durable expression of an encoded cargo [5, 6]. saRNA holds promise as a platform capable of addressing the shortcomings of mRNA by decreasing the dose of vaccines and administration frequency of protein-encoding therapeutics. Decreasing doses will mitigate vaccine related side effects and reduce the occurrence of rare but serious adverse events [7–9]. Additionally, order-of-magnitude reductions in dose requirements will significantly bolster the manufacturing capacity to enhance production speed and democratize the distribution of vaccines against emerging pathogens. However, early clinical evidence from saRNA vaccine trials show decreased efficacy and reduced neutralizing antibody levels compared to mRNA [10]. The early and intense activation of the type I interferon response induced by saRNA detection hinders replication and antigen expression [5, 11, 12]. Strategies to decrease the immunogenicity of saRNA are urgently needed to enable further clinical development and translation.

Prior efforts to decrease the immunogenicity of saRNA focus largely on sequence evolution, co-expression of viral inhibitory proteins, and optimization of the delivery vehicle [13–15]. While useful in their respective applications, all prior approaches fail to achieve a universal method of mitigating the interferon response and improving saRNA expression. The best tools for such improvements are modNTPs. However, the current understanding of the field is that the incorporation of modNTPs into saRNA abrogates downstream efficacy, as reported by multiple independent groups [5, 16–20]. Thus, modified nucleotides are not currently viewed as a viable strategy to decrease immunogenicity.

A hypothesis for this observed incompatibility is the alteration of vital saRNA secondary structures [16, 21]. saRNAs traditionally utilize alphavirus sequences, replacing the structural genes with the genes encoding the cargo(s) of interest [22]. The resulting synthetic construct encodes both an RdRp and the cargo(s) on the same RNA strand. Once expressed, the RdRp recognizes conserved sequence elements (CSEs) at the 5’ and 3’ ends to enable transcription of full-length copies of the entire saRNA construct. Later in its lifecycle, the RdRp complex also recognizes a sub-genomic promoter (SGP) to enable amplification of a truncated transcript encoding the cargo(s). The incorporation of modified nucleotides alters the stability or accessibility of specific base pairs, change hydrogen bonding patterns, and/or shift RNA hydrophobicity to stabilize or inhibit RNA-protein interactions [23–25]. Incorporating specific modNTPs, like N1mΨ may alter CSE or SGP structure and prevent effective RdRp recognition.

Here, we report the unexpected finding that several modified nucleotides at 100% substitution are compatible with saRNA and enhance its resulting *in vitro* and *in vivo* performance. A screen to identify modified modNTPs incorporated into a functional saRNA reporter system reveals two cytidine and one uridine modNTPs with activity greater than unmodified saRNA, while the vast majority of modNTPs, notably N1-methylpseudouridine (N1mΨ), yielded minimal transfection. Modified saRNA results in a decreased type I interferon response in primary human peripheral blood mononuclear cells (PBMCs) and increased reporter expression in several cell lines. Modified saRNA resulted in significantly longer expression of a luciferase reporter *in vivo* compared to modified mRNA. In a challenge study of SARS-CoV-2, the modified saRNA provides protection at significantly lower doses than modified mRNA and greater efficacy than unmodified saRNA.

## RESULTS

### Identification of modified nucleotides capable of maintaining saRNA activity

We synthesized a library of saRNA constructs through *in vitro* transcription, where all nucleotides were completely substituted with modified counterparts. To assess their functionality, these constructs encoded an mCherry reporter and were transfected into HEK293-T cells by cationic lipofection (Figure 1B). We identified three modNTPs that imparted significantly elevated transfection efficiencies compared to the N1-methylpseudouridine (N1mΨ) modified construct that resulted in minimal transfection. Flow cytometry analysis (Figure 1C, 1D) and live cell microscopy (Figure 1D) revealed that constructs with complete substitution of 5-hydroxymethylcytidine (5OHmC), 5-methylcytidine (5mC), or 5-methyluridine (5mU) exhibited transfection efficiencies 14-fold, 10-fold, and 8-fold higher, respectively, in comparison to the N1mΨ modified construct. The transfection efficiencies for the identified modNTPs were equal to or greater than the wildtype unmodified control (Figure S1A). Notably, in the HEK293-T cells transfected by lipofection for high-throughput screening, the mean cargo expression intensity was reduced compared to the wild-type construct for 5OHmC and 5mC (Figure S1B). Substitution with the identified modNTPs resulted in functional constructs when synthesized with Cap-0 structures (ARCA) or Cap-1 structures (CleanCap AU). However, a significant increase in expression intensity was observed when constructs contained a Cap-1 structure (Figure S1D).

**Figure 1:**
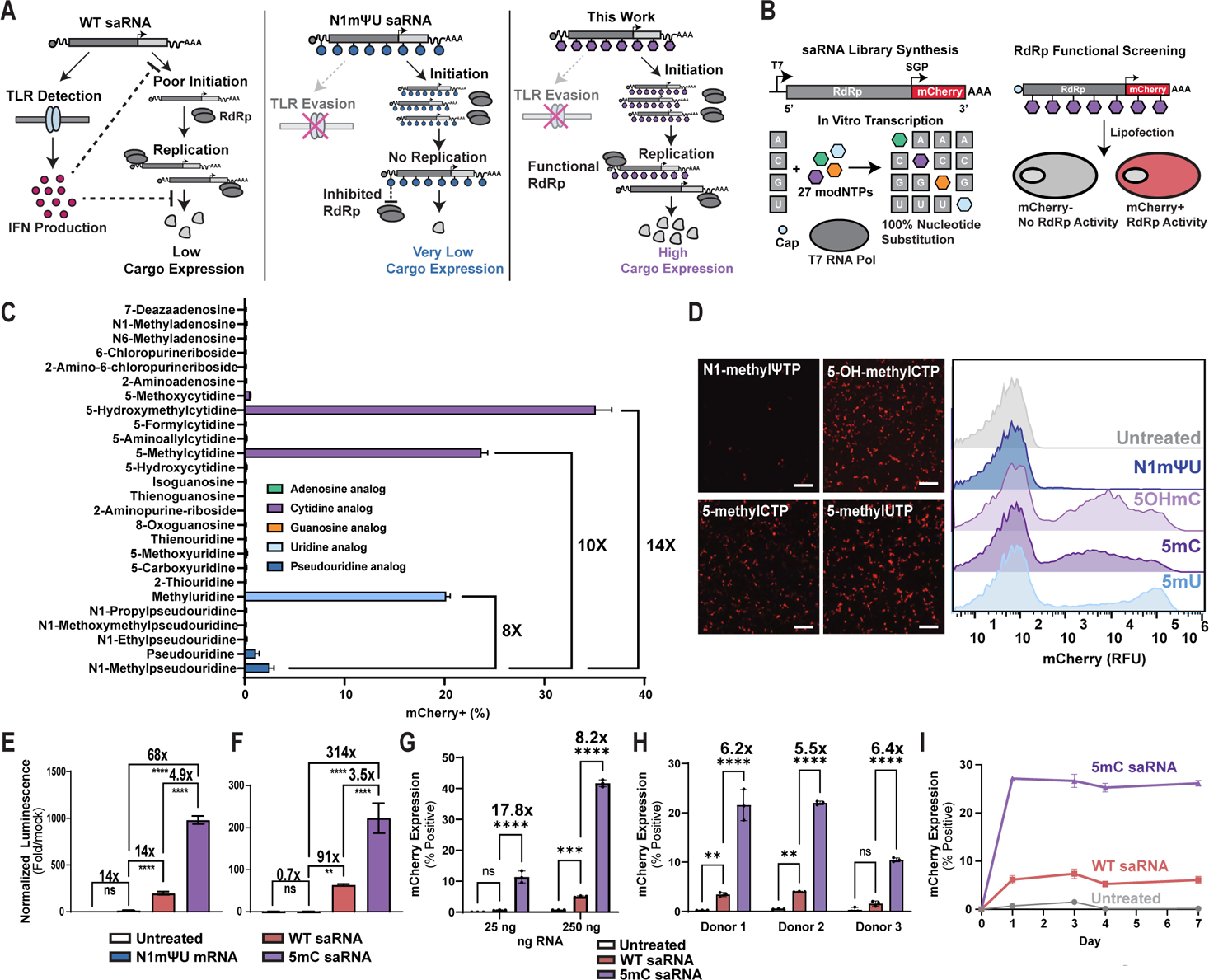
Identification of modified nucleotides compatible with self-amplifying RNA and their *in vitro* bioactivity. **A)** Schematic illustrating the limitations of unmodified saRNA, N1mΨ modified saRNA, and the advantages of saRNA with compatible modNTPs. **B)** Workflow for synthesizing a library of saRNA with complete substitution of modified nucleotides and transfecting the library encoding an mCherry reporter into HEK293-T cells using lipofection. **C)** Flow cytometry results measuring the percentage of HEK293-T cells transfected with the library of modified saRNA. Error bars represent the standard deviation of n = 3 biological replicates. **D)** Live-cell fluorescent microscopy images of selected modified nucleotides and representative flow cytometry histograms in HEK293-T cells. **E)** Expression levels 24 hours after transfection of 10 ng of RNA encoding a luciferase reporter in HEK293-T cells with SM102 LNPs. Luciferase signal is presented as fold-change compared to untransfected mock treated cells. Error bars represent the standard deviation of n = 4 biological replicates. **F)** Expression levels 24 hours after transfection of 10 ng of RNA encoding a luciferase reporter in C2C12 cells with SM102 LNPs. Luciferase signal is presented as fold-change compared to untransfected mock cells. Error bars represent the standard deviation of n = 4 biological replicates. **G**) Transfection efficiency 24 hours after LNP transfection with 25 ng and 250 ng of RNA encoding an mCherry reporter in Jurkat cells. Error bars represent the standard deviation of n = 3 biological replicates. **H)** Transfection efficiency 24 hours after LNP transfection of 500 ng of RNA encoding an mCherry reporter in primary human CD3+ T cells from three different donors. n = 3 biological replicates per group. **I)** Expression time course after LNP transfection with 100 ng of unmodified or 5mC modified saRNA in Jurkat cells. Error bars represent the standard deviation of n = 3 biological replicates. Two-way ANOVA with Tukey’s multiple comparisons correction, **** p < 0.0001, ** p < 0.01, * p < 0.05.

### Enhanced expression resulting from saRNA modification across diverse cell types *in vitro*

Motivated by the functionality of fully substituted saRNA, we sought to further investigate their activity *in vitro*. To assess the overall protein expression resulting from transfection with WT saRNA, 5mC saRNA, and N1mΨ mRNA, we employed a luciferase-based reporter construct. These constructs were loaded into lipid nanoparticles (LNPs). HEK293-T and C2C12 cells were transfected with LNPs containing 10 ng of RNA. Notably, the 5mC modified saRNA exhibited a remarkable 4.9-fold increase in expression over WT saRNA, corresponding to a substantial 68-fold increase in expression over the N1mΨ mRNA in HEK293-T cells (Figure 1E). In C2C12 cells, a 3.5-fold increase in expression was observed for the 5mC modified saRNA, resulting in a 314-fold increase compared to the N1mΨ mRNA, which resulted in limited expression at the low dosage administered (Figure 1F). A dose response experiment in HEK cells showed significantly improved expression at doses as low as 1 ng (Figure S2A). Next, we conducted transfection experiments with the modified saRNA in Jurkat T cells, a notoriously hard to transfect cell line by mRNA containing LNPs. To assess the expression profile at the individual cell level, we employed an mCherry reporter combined with flow cytometry analysis. A significant 17.8-fold improvement in transfection efficiency was observed at the lower dose of 25 ng and an 8.2-fold enhancement was observed at the higher dose of 250 ng (Figure 1G, Figure S2B, Figure S2C). A similar expression profile was observed when the luciferase constructs were transfected at 250 ng (Figure S2D). In a time course study, the expression was durable over 7 days from the modified saRNA in Jurkat cells (Figure 1I, Figure S2E). To assay the impact of fully substituted saRNA in primary cells, we treated CD3+ T cells derived from three different human donors with saRNA loaded-LNPs. All donors exhibited a 5-6-fold increase in transfection efficiency and significant expression of the mCherry reporter gene. This corresponded to approximately 1-3% transfection efficiency for the wild-type (WT) saRNA and approximately 10-20% for the 5mC-modified saRNA (Figure 1H, Figure S3A, Figure S3B). The expression of the mCherry reporter was detected for at least 5 days (Figure S3B).

### Reduced interferon response from modified saRNA in human PBMCs

To characterize the interferon response caused by wildtype or modified saRNA, we cultured human peripheral blood mononuclear cells (PBMCs) from three distinct donors with lipid nanoparticles (LNPs) loaded with saRNA (Figure 2B). Gene expression analysis revealed that saRNA treatment induces a significant increase in the expression of early interferon-related genes, namely IFN-α1, IFN-α2, and IFN-β1, after 6 hours (Figure 2C-2E). However, when 5OHmC or 5mC are incorporated, there was a large reduction in the expression of these early IFN genes. Specifically, IFN-α1 and IFN-β1 exhibited a reduction of more than 8.5-fold, while IFN-α2 showed a reduction of over 3-fold. N1mΨ further reduced expression of the IFN genes but resulted in decreased transfection efficiency and reporter expression (Figure 1C). The analysis of IFN-α subtypes in media from a single donor (Figure 2F) was consistent with the gene expression analysis. Additionally, a longitudinal analysis of human IFN-β in media indicated that modNTP saRNA effectively suppresses IFN-β expression. In all donors, no detectable expression of IFN-β was observed after 5mC and N1mΨ saRNA treatment, and only one donor exhibited detectable levels of IFN-β after 5OHmC saRNA treatment (Figure 2G).

**Figure 2.**
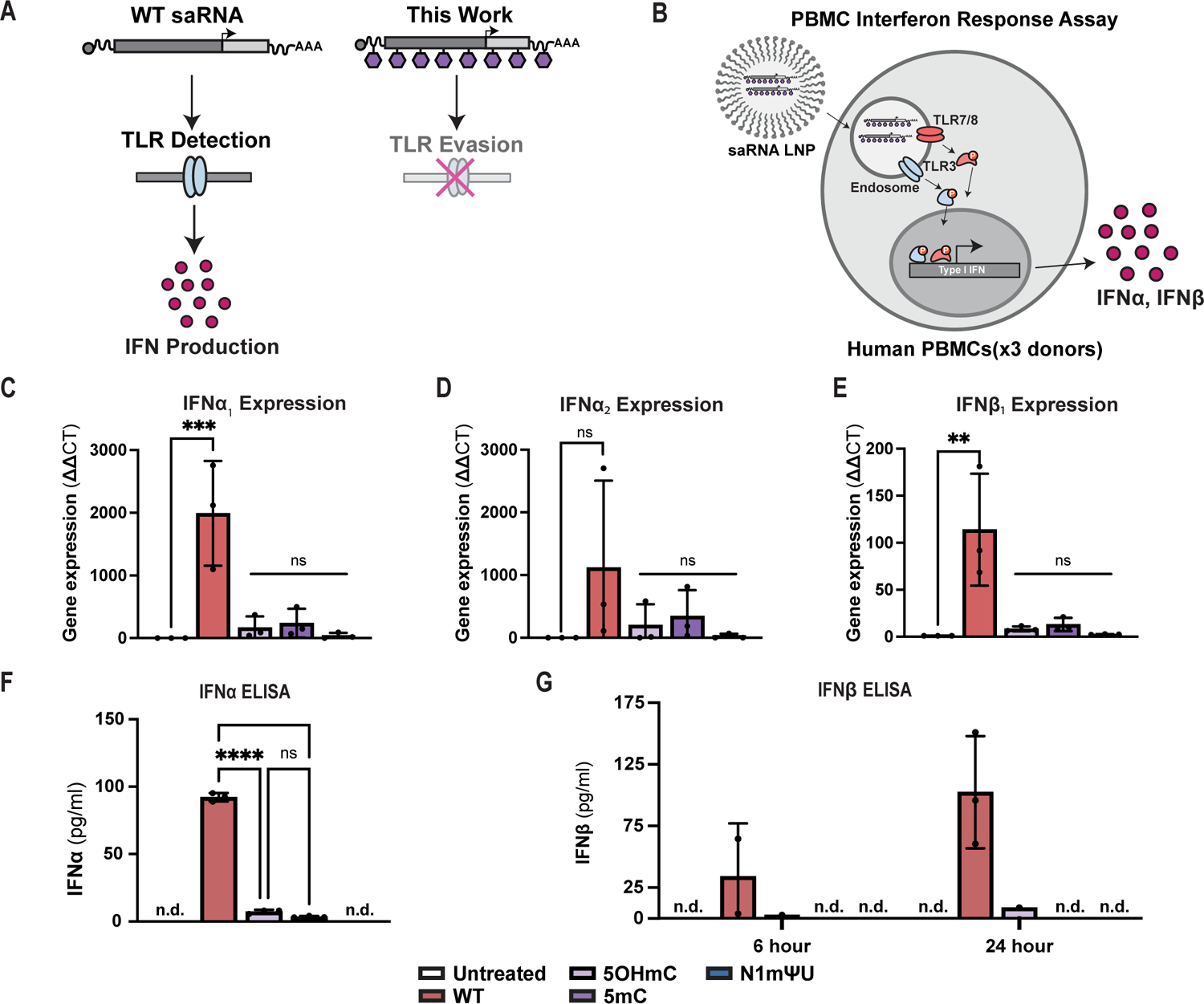
Evaluation of immunogenicity of modified saRNA in human PBMCs. **A)** Diagram depicting hypothesis of modified saRNA evading TLR detection, leading to reduced interferon production. **B)** Assay for detection of early interferon response from transfection of human PBMCs with unmodified or modified saRNA. Gene expression analysis of **C)** IFN-α_1_, **D)** IFN-α_2_, and **E)** IFN-β_1_ in RNA harvested from unique human PBMC donors (n=3) after 6-hour treatment with saRNA. **F**) IFN-α (all-subtype) protein level analysis by ELISA after 6 hours from a single PBMC donor treated with unmodified or modified saRNA with n = 3 biological replicates. **G**) 6 hour and 24-hour analysis of IFN-β protein levels by ELISA from a single PBMC donor treated with unmodified or modified saRNA with n = 3 biological replicates. Error bars represent standard deviation of n = 3 biological replicates. Statistical significance determined by ANOVA, controlling for multiple comparisons using Dunnett’s method. *** p < 0.001, **** p < 0.0001. n.d. not determined/below limit of detection.

### Development and validation of a low-dose modified saRNA SARS-CoV-2 vaccine

Next, we generated non-replicating mRNA (Spike mRNA) and self-amplifying RNA (Spike saRNA) encoding a K986P and V987P stabilized spike protein of SARS-CoV-2 derived from the Wuhan-1 strain (Figure 3A). The constructs were assembled by *in vitro* transcription. We transfected these RNAs into HEK and C2C12 cells via LNPs. In HEK cells, saRNA modified with 5OHmC, 5mC, and 5mU exhibited approximately twice the protein expression compared to wildtype saRNA, as assessed by flow cytometry (Figure 3B). Consistent with previous findings, N1mΨ modification of saRNA resulted in suppressed expression. In C2C12 cells, both 5OHmC and 5mC modifications significantly improved transfection efficiency, leading to approximately 2-fold higher protein expression than the unmodified saRNA construct and approximately 8-fold higher protein expression than N1mΨ modified non-replicating mRNA (Figure 3C, 3D, Figure S4A). These findings were further supported by ELISA analysis, which confirmed enhanced protein expression for the modified saRNA constructs (Figure S4B). Interestingly, the 5mU modification did not result in protein expression in C2C12 cells. Similar *in vitro* experiments were performed to evaluate the expression of the flu hemagglutinin (HA) antigen and an expression profile consistent with previous experiments was observed (Figure S4C+D).

**Figure 3.**
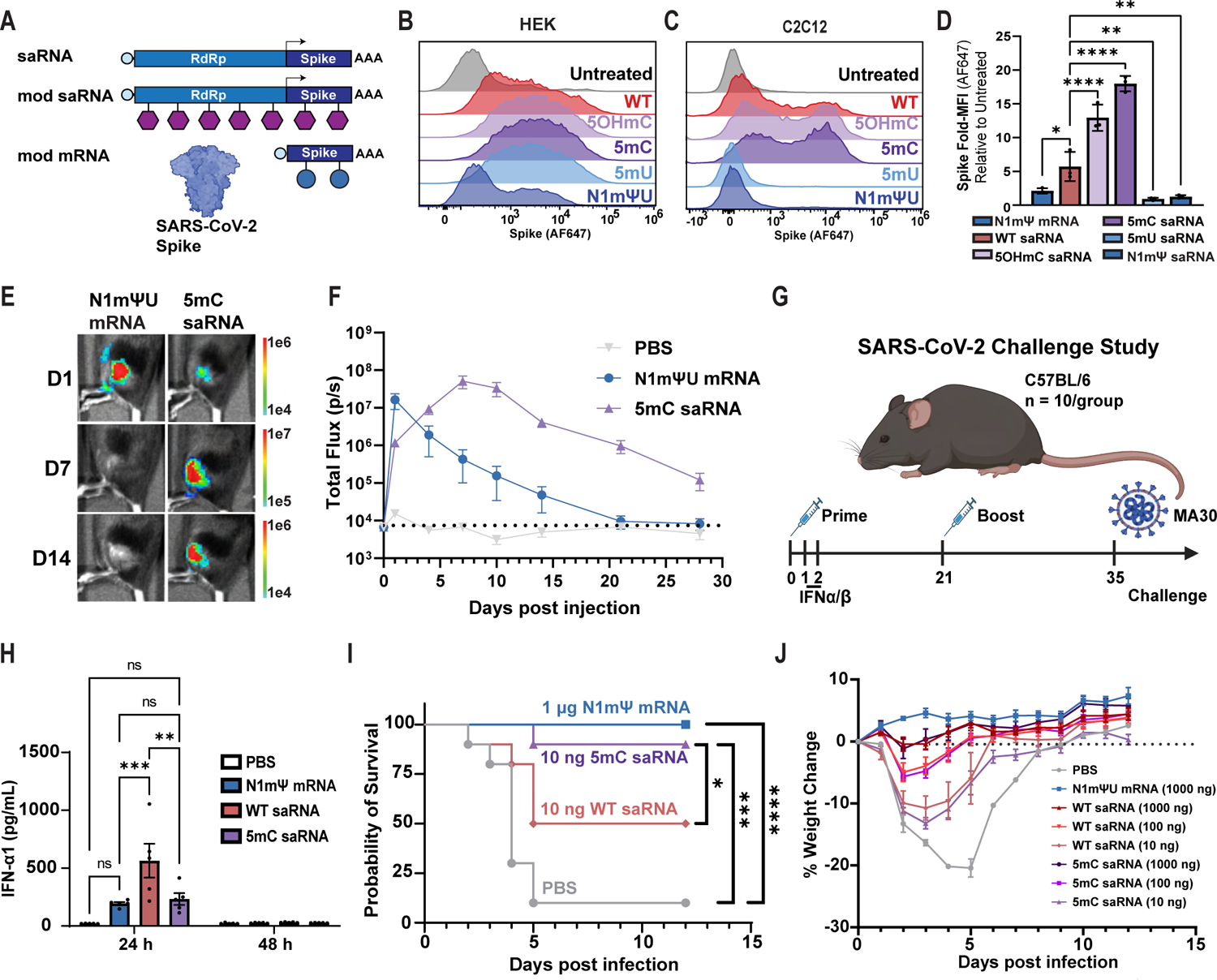
Development and characterization of a fully modified saRNA vaccine against SARS-CoV-2. **A**) Schematic illustrating the different RNA formats for expressing the SARS-CoV-2 Spike protein, compared *in vitro* and *in vivo*. This includes non-replicating N1mΨ mRNA, wildtype self-amplifying RNA, and 5mC modified self-amplifying RNA. **B**) Expression of Spike protein detected by flow cytometry 24 hours after LNP transfection with 100 ng of RNA in HEK293-T cells with n = 3 biological replicates. **C**) Expression of Spike protein detected by flow cytometry 24 hours after LNP transfection with 100 ng of RNA in C2C12 cells with n = 3 biological replicates. **D**) Median fluorescence intensity (MFI) of anti-Spike AF647 staining in C2C12 cells. MFI is relative to cells that were not transfected but were stained with anti-Spike AF647. Error bars represent the standard deviation of n = 3 biological replicates. **E**) Bioluminescent images of mice at different time points after i.m. injection with LNPs containing 2.5 µg of luciferase encoding N1mΨ mRNA (left) or 5mC saRNA (right). Scale bar indicates radiance. **F)** Total flux from BLI imaging of mice injected intramuscularly with LNPs containing 2.5 µg of luciferase encoding N1mΨ mRNA or 5mC saRNA (n = 5 biological replicates). Dashed line indicates average signal for the PBS group over the duration of the study. Error bars indicate standard error mean. **G)** Study design for the SARS-CoV-2 challenge study in C57BL/6 mice. Mice were vaccinated in a prime-boost scheme, serum was collected to analyze interferon response, and mice were challenged with MA30 on day 35. **H**) IFN-α1 expression in serum collected 24 hours or 48 hours after initial vaccination with 1000 ng of RNA in LNPs. Error bars represent the standard error mean for n = 5 biological replicates. **I**) Survival of mice after challenge with a lethal challenge of MA30 virus. n = 10 mice per group. **J**) Weight change of mice after challenge with mouse-adapted SARS-CoV-2 MA30 virus. Error bars indicate standard error mean. Statistical significance determined by ANOVA, controlling for multiple comparison’s using Dunnett’s method. ** p < 0.005, *** p < 0.001, **** p < 0.0001. For survival study statistics, a log-rank (Matel-Cox) test was used between groups. **** = p < 0.0001, *** = p < 0.001.

Next, the expression levels and kinetics of modified saRNA and modified mRNA were compared *in vivo* via bioluminescent imaging (BLI). C57BL/6 mice (n = 5 per group) were intramuscularly (i.m.) administered PBS, 2.5 µg of N1mΨ modified non-replicating mRNA or 5mC modified saRNA encoding firefly luciferase. The mRNA group exhibited a greater total flux at 24 hours (Figure 3E). In contrast, the 5mC saRNA group showed a consistent increase in signal, reaching peak expression 7 days after the initial injection, which was four times greater than that of the mRNA group (Figure 3F). Notably, the modified saRNA group displayed significantly prolonged expression, maintaining an average signal above the background observed in the PBS treated group throughout the entire 28-day study duration (Figure 3F).

Motivated by the successful antigen expression observed *in vitro* and durable expression *in vivo,* we sought to evaluate the effectiveness of a modified saRNA vaccine against SARS-CoV-2 infection. C57BL/6 mice (5 male and 5 female) were intramuscularly (i.m.) administered 10 ng, 100 ng, and 1000 ng of WT Spike saRNA or 5mC Spike and boosted at 35 days post initial vaccination (Figure 3E). As positive and negative controls, additional groups received 1000 ng of N1mΨ non-replicating mRNA or vehicle (PBS), respectively. To assess the early interferon response in mice, serum was collected from mice vaccinated with the 1000 ng dose at 24 hours and 48 hours. The mice vaccinated with WT saRNA exhibited significantly increased levels of serum IFN-α1 compared to N1mΨ or 5mC saRNA vaccinated mice. Conversely, mice receiving N1mΨ mRNA or 5mC saRNA displayed significantly reduced levels of serum IFN-α1, indicating a decrease in TLR signaling attributable to the modified nucleotides. By 48 hours post vaccination, IFN-α1 was no longer detectable in the serum of any mice (Figure 3G). Across all samples, IFN-β was not detectable at 24 hours or 48 hours (Figure S6A). On day 35, 14 days after the final vaccination, the mice were infected intranasally with 1×10^5^ PFU of mouse adapted SARS-CoV-2 (MA30) [26]. The mice vaccinated with either WT Spike saRNA or 5mC Spike saRNA at 100 ng and 1000 ng showed 100% survival and minimal weight loss (Figure S6B, Figure 3I). However, mice receiving the 10 ng dose of WT Spike saRNA showed severe weight loss and 50% lethality. In contrast, mice vaccinated with the 10 ng dose of 5mC demonstrated only 10% lethality, a statistically significant improvement in survival compared to WT Spike saRNA (Figure 3I+J).

## DISCUSSION

As a technology, self-amplifying RNA promises lower dose vaccines and off-the-shelf, *in situ*, long-lasting, and non-integrating cell and gene therapies. Unfortunately, the clinical reality thus far paints a less encouraging picture [6]. Despite success in preclinical models, saRNA vaccines show reduced seroconversion in human trials [10]. An analysis of gene expression in vaccinated humans reveals sharp activation of the innate immune system [27]. Preclinical and clinical data also confirm that the early and intense interferon response inhibits antigen expression [5, 11, 12, 28]. A mechanism to evade the early interferon response and improve saRNA transfection will unlock the true potential of the platform.

Prior strategies to enhance the efficacy of saRNA focus on reducing the innate immune response by optimizing the sequence, encoded cargo, and delivery vehicle. Evolved RdRp variants capable of decreasing the interferon response enhance the anti-tumor efficacy of an IL2 encoding saRNA [13]. While promising, it is unclear how evolution of the RdRp sequence will affect performance in contexts beyond intratumoral immunotherapy. Co-expression of viral proteins that block the interferon response results in improved titers in a vaccine against RABV [14]. It is anticipated that this strategy will enhance activity only in cell types lacking robust TLR sensing that are amenable to saRNA transfection. In the context of vaccination, the delayed interferon response resulting from cytosolic sensors likely has an adjunctive effect. For therapeutics, it may be desirable for saRNA constructs to evade TLR detection and to inhibit the downstream interferon response triggered by cytosolic sensors. Recently, a localizing cationic nanocarrier formulation (LION) limits the delivery of saRNA to the injection site and thereby decreases the systemic type I interferon response [15]. Dependency on the cationic delivery vehicle may pose challenges in achieving efficient delivery to specific tissue or cell types, potentially requiring higher doses, and thereby imposing constraints on its broader therapeutic applications. In sum, prior research demonstrates need for a universal approach to control the interferon response and improve saRNA translation.

The best understood tool for suppression of the interferon response in non-replicating RNA therapeutics is the incorporation of modified nucleotides, as exemplified by N1mΨ modified mRNA. The current understanding in the field is that modified nucleotides (modNTPs) are incompatible with saRNA and that saRNA performance decreases proportional to incorporation percentage. In fact, multiple studies report the failure of modified saRNA, since the first discovery of the impact of modified nucleotides on mRNA [5, 16–20, 29–33]. We hypothesized that modified nucleotides compatible with saRNA exist, considering the numerous RNA modifications observed in eukaryotic cells and their ubiquitous presence in RNA viruses [34].

Here, we provide the first evidence of biologically active, fully substituted self-amplifying RNA containing 5OHmC, 5mC, or 5mU. These three modNTPs maintain saRNA activity at levels equal to or greater than an unmodified construct (Figure 1). These identified modified nucleotides significantly enhance transfection efficiency in HEK293-T, C2C12, and Jurkat cells (Figure 1). This potentiation translates to transgenes in T cells (Figure 1) and vaccine antigens in muscle cells (Figure 3, Figure S4). Importantly, significantly reduced expression of type I interferons occurs in transfected human PBMCs *in vitro* (Figure 2) and systemically *in vivo* (Figure 3). The significant reduction in serum IFN-α levels offers compelling evidence of diminished systemic immunogenicity, a factor frequently acknowledged as the primary remaining limitation associated with saRNA. The expression durability of modified saRNA is significantly greater than modified mRNA (Figure 3). Extensive *in vitro* and *in vivo* validation of a modNTP substituted saRNA vaccine candidate provides proof-of-concept for enhanced clinical efficacy. In a lethal challenge study, just 10 ng of modNTP substituted saRNA performs statistically better than unmodified saRNA, truly enabling the claims of saRNA as an ultra-low dose vaccine platform (Figure 3). Early studies presented in this work suggest modified self-amplifying RNA as a platform technology capable of revolutionizing modern RNA therapeutics.

Concurrent with our discovery, a team lead by Akahata reported a week earlier that incorporation of 5mC in self-amplifying RNA affords a functional construct as a vaccine against SARS-CoV-2 [35]. No *in vitro* preclinical data or comparisons to unmodified saRNA or modified mRNA were provided. The group conducted a phase 1 human safety study of a 5mC modified saRNA SARS-CoV-2 booster with healthy patients in Japan. Patients were vaccinated with 1, 3, 7.5, and 15 ug of LNP doses of 5mC modified saRNA and IgG titers at day 28 increased on average ≍ 3-fold for all doses, with no comparative arms to modified mRNA or unmodified saRNA. Importantly this work highlights the potential clinical utility of modified saRNA via positive human safety studies.

The discovery of modNTPs capable of maintaining saRNA functionality is a major step towards a deeper understanding of alphavirus transcriptional activity and control. Results here demonstrate that RdRp transcriptional activity is maintained with the incorporation of specific modNTPs but lost with gold standard modNTPs such as N1mΨ. The observation of launch from fully modified saRNA means the two events must still be occurring: 1) cap-dependent translation of the RdRp and 2) the RdRp continues to recognize at least some of the conserved elements required for negative strand synthesis. Negative strand synthesis occurs intracellularly, and as a result, only wild type nucleotides will be incorporated into the new negative strand. Taking this and the results from N1mΨ saRNA together suggest that modNTPs primarily impact RdRp-saRNA interaction, and not the initial cap-dependent translation event. This is just the first step towards a deeper mechanistic understanding of saRNA and this work provides motivation to pursue basic RNA biology studies with modified RNA. This work begs numerous questions, such as the reason behind the exclusive saRNA activity of purine analogs and the complete elimination of activity by all pyrimidine analogs. Additionally, the compatibility of 5-methyluridine in HEK293-T cells but not in C2C12 cells suggests a potential epitranscriptomic discrepancy and mechanism of cell-type specificity. Explorations into the effect of modified nucleotides in the broader family of alphaviruses may shed light on these remaining questions.

Other emerging RNA formats, notably circular RNA, may benefit from similar endeavors to screen for compatible modified nucleotides [36–39]. Circular RNAs utilize IRES sequences to drive translation, and these sequences are commonly derived from RNA and DNA viruses. To date, modified nucleotides compatible with circular RNA and IRES sequences remain elusive. Performing similar screens with circular RNA may shed light on the compatibility and interactions of modified nucleotides with diverse RNA structures.

In conclusion, this work presents a significant advancement to the field of self-amplifying RNA, and RNA therapeutics more broadly. This is the first known embodiment of fully substituted self-amplifying RNA with enhanced function, which goes against the dogmatic thinking of the field [40]. Modified nucleotides overcome the major barrier to successful translation of saRNA into the clinic, which is the early interferon response preventing launch and production of cargo proteins. This discovery considerably broadens the potential scope of self-amplifying RNA, enabling entry into previously impossible cell types, as well as the potential to apply saRNA technology to non-vaccine modalities such as cell therapy and protein replacement.

## MATERIALS AND METHODS

### Template design and synthesis

All mRNA and saRNA templates took the form of linearized plasmid DNA. All saRNA templates were generated from a plasmid encoding NSP1-4 of the Venezuela Equine Encephalitis Virus (VEEV) under either a CleanCap AU (TriLink BioTechnologies) compatible T7 promoter (Promoter seq: TAATACGACTCACTATAGAT) or a GTP/ARCA compatible T7 promoter (Promoter seq: TAATACGACTCACTATAGGAT. Sequences from VEEV were derived from T7-VEE-GFP, a gift from Steven Dowdy (Addgene plasmid #58977; http://n2t.net/addgene:58977; RRID: Addgene_58977) [41]. Reporter plasmids were generated by inserting the coding sequences for mCherry or firefly luciferase between 5’ and 3’ UTR sequences derived from human beta-globin. Antigen sequences are as follows: Influenza A Hemagglutinin (Influenza A virus (A/California/07/2009(H1N1)), NCBI Ref Seq: YP_009118626.1), SARS-CoV-2 Spike (Wuhan-Hu-1, NCBI Ref Seq: YP_009724390.1) with K986P and V987P stabilizing mutations. Plasmids were cloned in DH5α *e coli.* Plasmids were purified using ZymoPURE™ II Plasmid Midiprep Kit (Zymo Research), linearized with MluI-HF for 3 hours at 37°C, and purified using QIAquick PCR Purification Kit (Qiagen).

### RNA synthesis

mRNA and saRNA was synthesized using MEGAscript T7 Transcription kit (Invitrogen™, ThermoFisher Scientific) with 1μg template and co-transcriptional capping using CleanCap AU (TriLink BioTechnologies). For IVT with ARCA, a final concentration of 12mM ARCA and 4mM GTP was used. For ARCA experiments were a GTP analog was utilized, the modNTP was substituted completely to a final concentration of 4mM. The modNTPs in all CleanCapAU IVT reactions were prepared according to manufacturer’s protocol. IVT for 3 hours at 37C was followed by 10min DNase treatment and 30 min post-transcriptional poly-adenylation using a Poly(A) Tailing Kit (Invitrogen™, ThermoFisher Scientific). saRNA was purified using MEGAclear™ Transcription Clean-up Kit (Invitrogen™ ThermoFisher Scientific) and eluted in DNase/RNase free H_2_O prior to storing at −80C. The A260 of an equimolar, 1mM mixture of all combinations of modNTPs and NTPs was empirically determined using a NanoDrop2000 (ThermoFisher Scientific). The adjusted factor for each modNTP was used to calculate RNA concentration to ensure consistent delivery between modNTPs (Table 1).

### modNTP saRNA screening in HEK293T

HEK293-T cells were grown in DMEM + 1mM sodium pyruvate + 10% FBS + 1% PS, plated at 70,000cells/cm^2^ in 96-well plates, and allowed to adhere overnight. mCherry saRNA was transfected using MessengerMax according to manufacturer protocols (Invitrogen™, ThermoFisher Scientific). 24 hours later, cells were prepared for flow cytometry analysis of mCherry expression. Cells were washed 1X in PBS, treated with 1X PBS +2mM EDTA for 5 mins, and then resuspended in FACS Buffer (1X PBS + 2% BSA). Data was acquired on the Attune NxT (Thermo Fisher Scientific) and analyzed on FlowJo (version 10.8.1) Fluorescence microscopy of selected modNTPs was performed on a Biotek Cytation 5 microscope prior to preparation for flow cytometry analysis.

### LNP Formulation

saRNA were encapsulated inside of LNPs with the following composition (mole percent): 50% (8-[(2-hydroxyethyl)[6-oxo-6-(undecyloxy)hexyl]amino]-octanoic acid, 1-octylnonyl ester [SM-102, Cayman]), 1.5% (1,2-dimyristoyl-rac-glycero-3-methoxypolyethylene glycol-2000 [DMG-PEG2K, Avanti]), 10% (1 1,2-dioleoyl-sn-glycero-3-phosphoethanolamine [DOPE, Avanti]), 38.5% cholesterol (Avanti). RNA was loaded at an N:P of 10. Prior to formulation, aqueous and lipid phases were separately sterile filtered through 0.22μm filters. Formulations were performed in sterile RNAse free conditions. After formulation, LNPs were dialyzed against sterile RNAse free 1X PBS for 24 hours at 4°C. LNP morphology was characterized via DLS using a NanoBrook Omni (Brookhaven Instruments). Encapsulation efficiency was determined using the QuantiFluor RNA System (Promega). To transfect primary T cells, mAb-targeted LNPs were formulated via post-insertion of DSPE-PEG2K-maleimide coupled to anti-CD3 reduced with TCEP (Clone OKT3, BioXCell).

### Expression Potentiation Assays

HEK and C2C12 cells were grown in DMEM + 10% FBS + 1% PS and were passaged every three days. 24 hours prior to transfection, the cells were washed with PBS, trypsinized, and plated in 200 µL DMEM at 50,000 cells/well for HEK and 25,000/well for C2C12. For transfection, LNPs containing mRNA or saRNA encoding luciferase were added dropwise to the cells in triplicate at 10 ng or 100 ng/well. After 24 hours, the luciferase expression was assayed using the BrightGlo system (Promega).

Jurkat cells were grown in RPMI + 10% FBS + 1% PS and maintained between 5×10^5^ and 1×10^6^ cells/ml. Prior to transfection, the cells were washed with PBS and plated in 200 µL RPMI at 250,000 cells/well. For transfection, LNPs containing saRNA with or without modified nucleosides were added dropwise to cells in triplicate at 25 or 250 ng/well. After 24 hours, a portion of the cells were harvested for flow analysis. The remainder of the cells were maintained in culture and assayed by flow at additional time points.

Primary human CD3+ T cells from 3 unique donors were isolated by negative selection (T Cell Enrichment Cocktail, Stemcell Technologies) from peripheral blood and were grown in RPMI + 10% FBS + 1% PS supplemented with 50 U/mL of IL2, 10 ng/mL IL7, 10 ng/mL IL15. To activate the cells, CD3/CD28 Dynabeads were added to the cells at a 1:1 bead to cell ratio for 24 hours. After activation, the Dynabeads were removed, and the T cells were rested for 24 hours before transfection. Primary T cells were washed with PBS and plated at 100K/well in RPMI + 10% FBS supplemented with 50 U/mL IL2. For transfection, anti-CD3 conjugated LNPs were dosed in triplicate at 500 ng/well in 200 µL.

### PBMC early interferon response

Human Peripheral Blood Mononuclear Cells (PBMCs) were thawed and rested for 24 hours in RPMI + 10% FBS + 1% PS. 500,000 cells (1M/ml) were plated in a 24 well plate. 250 ng of saRNA encapsulated in LNPs, formulated as previously described, was added dropwise to the PBMCs. After 6 or 24 hours, media was harvested by centrifugation and analyzed for interferon expression by ELISA. (Human IFN-alpha All Subtype Quantikine ELISA - R&D Systems) (Human IFN-beta DuoSet ELISA R&D Systems). cDNA was generated using High-Capacity cDNA Reverse Transcription Kit (Applied Biosystems), according to manufacturer’s protocols. qPCR for specific genes was carried out using TaqMan probes in TaqMan advanced master mix (Thermo Fisher Scientific). UBC-VIC was multiplexed as the endogenous control for all qPCRs.

### Antigen production in C2C12 Cells

C2C12 cells were grown in DMEM + 10% FBS + 1% PS, plated at 25,000 cells/well in 96-well TC treated plates, and allowed to adhere overnight. For transfection, LNPs containing mRNA or saRNA encoding for each antigen were added to cells in triplicate at 100 ng/well, except for HA which was dosed at 25 ng/well. After 24 hours cells were harvested for flow analysis using the following mAbs: SARS-CoV-2 Spike Protein (RBD) Human monoclonal (eBiosciences - Clone: P05Dhu – AF647 conjugated), Influenza A H1N1 (A/California/07/2009) Hemagglutinin, Rabbit polycloncal: (1:1000 dilution) (SinoBiological - Cat: 11085-T62 – APC conjugated). Identical treatments were performed for ELISA confirmation. Cells for ELISA were washed twice in PBS and then resuspended in 1X PBS + 0.1% triton X-100. Cells were freeze-thawed once before thorough pipette mixing to ensure lysis and membrane disruption. ELISA was performed with antigen specific sandwich ELISA kits according to manufacturer’s protocol (SinoBiological).

### Institutional approvals

All animal experiments described in this study were performed in accordance with protocols that were reviewed and approved by the Institutional Animal Care and Use and Committee of Boston University (PROTO202100000026, PROTO202000020, and PROTO201800600). All mice were maintained in facilities accredited by the Association for the Assessment and Accreditation of Laboratory Animal Care (AAALAC). Vaccination studies and replication-competent SARS-CoV-2 experiments were performed in a biosafety level 2 and 3 laboratory (BSL-3), respectively at the Boston University National Emerging Infectious Diseases Laboratories (NEIDL). Bioluminescent imaging experiments were performed at Boston University.

### Bioluminescent imaging

Mice were injected intramuscularly (i.m.) in the left hind limb with 50 µL of PBS or LNPs containing 2.5 µg of mRNA or saRNA. For imaging, the mice were anesthetized with 1-2% isoflurane and were injected intraperitoneally (i.p.) with D-luciferin 10 minutes prior to imaging.

### Virus production cell culture

VeroE6 cells were grown in Dulbecco’s modified Eagle’s medium (DMEM) (ThermoScientific, Waltham, MA, USA) supplemented with 10% (v/v) heat-inactivated fetal bovine serum (FBS) (Bio-Techne, R&D systems, Minneapolis, MN, USA) and 1% (v/v) penicillin streptomycin (P/S) (ThermoScientific, Waltham, MA, USA). A549-hACE2/hTMPRSS2 cell lines were maintained in DMEM containing 10% FBS and 1% PS supplemented with 2.5 ug/mL Puromycin and Blasticidin. All cell lines were maintained in a cell incubator at 37°C with 5% CO_2_.

### SARS-CoV-2 isolate stock

All replication-competent SARS-CoV-2 experiments were performed in a BSL-3 facility at the Boston University NEIDL. The clinical isolate named 2019-nCoV/USA-WA1/2020 strain (NCBI accession number: MN985325) of SARS-CoV-2 was obtained from BEI Resources (Manassas, VA, USA). SARS-CoV-2 MA30 was generously provided by Dr. Stanley Perlman (University of Iowa). SARS-CoV-2 Delta variant was generously provided by Dr. John Connor (Boston University). Viral stocks were prepared and titered as previously described [42].

### Mouse vaccine studies

8-10 weeks old male and female C57BL/6J mice were obtained from Jackson Laboratories (Bar Harbor, ME, USA, Catalog # 000664). In the NEIDL BSL-2 and −3 facility, mice were group-housed by sex in Techniplast green line individually ventilated cages (Techniplast, Buguggiate, Italy). The room was maintained with a 12:12 light cycle at 30-70% humidity. Mice were injected intramuscularly (i.m.) in the hind limb with 50 µL of PBS or LNPs containing 10 ng, 100 ng, or 1000 ng of mRNA or saRNA. At 24- and 48-hours post vaccination, serum was collected via submandibular bleeding from groups of five mice from the 1000 ng vaccinated groups for analysis of interferon response. At 28 days post vaccination, mice were administrated i.m. with a booster dose of PBS or vaccine (50 µL; similar dosage as primary vaccination). At day 35 post vaccination, mice were transferred into BSL-3 and challenged with MA30 virus.

### SARS-CoV-2 challenge experiments

Vaccinated and boosted mice were intranasally inoculated with 1×10^5^ PFU SARS-CoV-2 MA30 virus resuspended in 50 µL of 1x PBS. Mice were inoculated under 1-3% isoflurane anesthesia. Infected mice were monitored and clinically scored for changes in weight, respiration, appearance, responsiveness and behavior over the course of 14-days post infection. Mice with a cumulative clinical score of 4 or 25% weight loss were euthanized.

### Serum preparation

Blood was collected at the designated time points via submandibular bleeding and serum was isolated by centrifuging blood in a benchtop centrifuge at 3500 RPM for 10 minutes at room temperature. Serum was then collected, transferred into a new Eppendorf tube and stored at −80°C for downstream analysis.

### Serum interferon analysis

Serum interferon alpha and beta were analyzed by ELISA (LEGEND MAX™ Mouse IFN-β ELISA Kit – BioLegend) (ELISA MAX™ Deluxe Set Mouse IFN-α1 – BioLegend).

## ACKNOWLEDGEMENTS

W.W.W. acknowledges support from the NIH U01CA265713, R01EB029483, R01DK132576, R01EB031904. M.W.G. acknowledges support from NIH U01CA265713 and a William Fairfield Warren Professorship. F.D. received support from Boston University startup funds. D.K. received support from an NIH T32 Immunology Training Program (T32AI007309). We thank the NEIDL animal core and NEIDL operation staff for their outstanding support. T.S. received support from a Lung Cancer Research Fellowship. J.E.M. received support from NSF GRFP. J.E.M. and J.R.K. were supported by an NIH T32 Translational Research in Biomaterials Training Program (T32 EB006359). This material is based upon work supported by the National Science Foundation Graduate Research Fellowship Program under Grant No. 2234657. Any opinions, findings, and conclusions or recommendations expressed in this material are those of the author(s) and do not necessarily reflect the views of the National Science Foundation. This research was supported in part by the National Institutes of Health training grant at Boston University, T32 EB006359. The content is solely the responsibility of the authors and does not necessarily represent the official views of the National Institutes of Health.

## DATA AVAILABILITY

Source data used for figure generation and statistical analysis are provided with the manuscript.

## CONTRIBUTIONS

J.E.M., J.R.K., M.W.G., and W.W.W. were involved in conceptualization. J.E.M. and J.R.K. designed and assembled RNA constructs and conducted *in vitro* experiments. D.K. and E.C. performed *in vivo* experiments involving the SARS-CoV-2 vaccine. T.S. and J.E.M. performed *in vivo* bioluminescent imaging. J.E.M., J.R.K., M.W.G, W.W.W., F.D. were involved in study design. All authors provided feedback and contributed to manuscript preparation.

