## Supplementary Information for "Complete substitution with modified nucleotides suppresses the early interferon response and increases the potency of self-amplifying RNA"

| **Chemical Name** | **Base Analog** | **Mol Weight** (g/mol) | **Abs** (260nm) | **Factor** |
|---|---|---|---|---|
| 7-Deazaadenosine | Adenosine | 506.1 | 5.04 | 55.976 |
| N1-Methyladenosine | Adenosine | 521.2 | 15.369 | 39.399 |
| N6-Methyladenosine | Adenosine | 521.2 | 14.777 | 39.727 |
| 6-Chloropurineriboside | Adenosine | 526.6 | 17.481 | 38.430 |
| 2-Amino-6-chloropurineriboside | Adenosine | 541.6 | 11.482 | 42.280 |
| 2-Aminoadenosine | Adenosine | 522.2 | 14.523 | 39.880 |
| Thienocytidine | Cytidine | 539.2 | *358.295 | n.d. |
| 5-Methoxycytidine | Cytidine | 513.2 | 13.912 | 34.580 |
| 5-Hydroxymethylcytidine | Cytidine | 513.1 | 14.618 | 34.140 |
| 5-Formylcytidine | Cytidine | 511.1 | 14.938 | 33.946 |
| 5-Aminoallylcytidine | Cytidine | 538.2 | 15.888 | 33.546 |
| 5-Methylcytidine | Cytidine | 497.1 | 15.01 | 33.847 |
| 5-Hydroxycytidine | Cytidine | 499.1 | 14.513 | 36.431 |
| Isoguanosine | Guanosine | 523.1 | 13.135 | 39.916 |
| Thienoguanosine | Guanosine | 539.2 | 11.774 | 41.100 |
| 2-Aminopurine-riboside | Guanosine | 507.2 | 10.584 | 42.112 |
| 8-Oxoguanosine | Guanosine | 539.1 | 13.191 | 39.951 |
| 5-Carboxymethylesteruridine | Uridine | 542.1 | 23.333 | 33.266 |
| Thienouridine | Uridine | 542.2 | 15.559 | 36.017 |
| 5-Methoxyuridine | Uridine | 514.1 | 13.408 | 37.210 |
| 5-Carboxyuridine+A17 | Uridine | 528.1 | 16.619 | 35.437 |
| 2-Thiouridine | Uridine | 500.2 | 14.605 | 36.376 |
| Methyluridine | Uridine | 498.1 | 15.047 | 36.115 |
| N1-Propylpseudouridine | Uridine | 526.2 | 13.523 | 37.186 |
| N1-Methoxymethylpseudouridine | Pseudouridine | 528.2 | 14.041 | 36.847 |
| N1-Ethylpseudouridine | Pseudouridine | 512.2 | 13.988 | 36.810 |
| N1-Methylpseudouridine | Pseudouridine | 498.1 | 14.557 | 36.396 |
| Pseudouridine | Pseudouridine | 484.1 | 14.861 | 36.160 |
| Wild type | - | | 13.865 | 38.288 |

**Table 1**.) Empirical determination of modified nucleotide absorbance conversion factors. Due to sample evaporation, as determined by significant increase in viscosity, thienocytidine concentration was hypothesized to be >100mM as listed. A wild-type absorbance factor was applied to thienocytidine A260. N.d. not determined.

| Group 1 | Group 2 | P value |
|---|---|---|
| 10 ng WT saRNA | 10 ng 5mC saRNA | 0.0497 |
| PBS | 10 ng 5mC saRNA | 0.0001 |
| 10 ng 5mC saRNA | 1000 ng N1mΨ mRNA | 0.3173 |
| PBS | 1000 ng N1mΨ mRNA | <0.0001 |
| PBS | 100 ng WT saRNA | <0.0001 |
| PBS | 100 ng 5mC saRNA | <0.0001 |
| PBS | 1000 ng WT saRNA | <0.0001 |
| PBS | 1000 ng 5mC saRNA | <0.0001 |

**Table 2**.) Summary of survival study statistics for group comparisons in Figure 3 and Figure S5. Log-rank (Mantel-Cox) test.

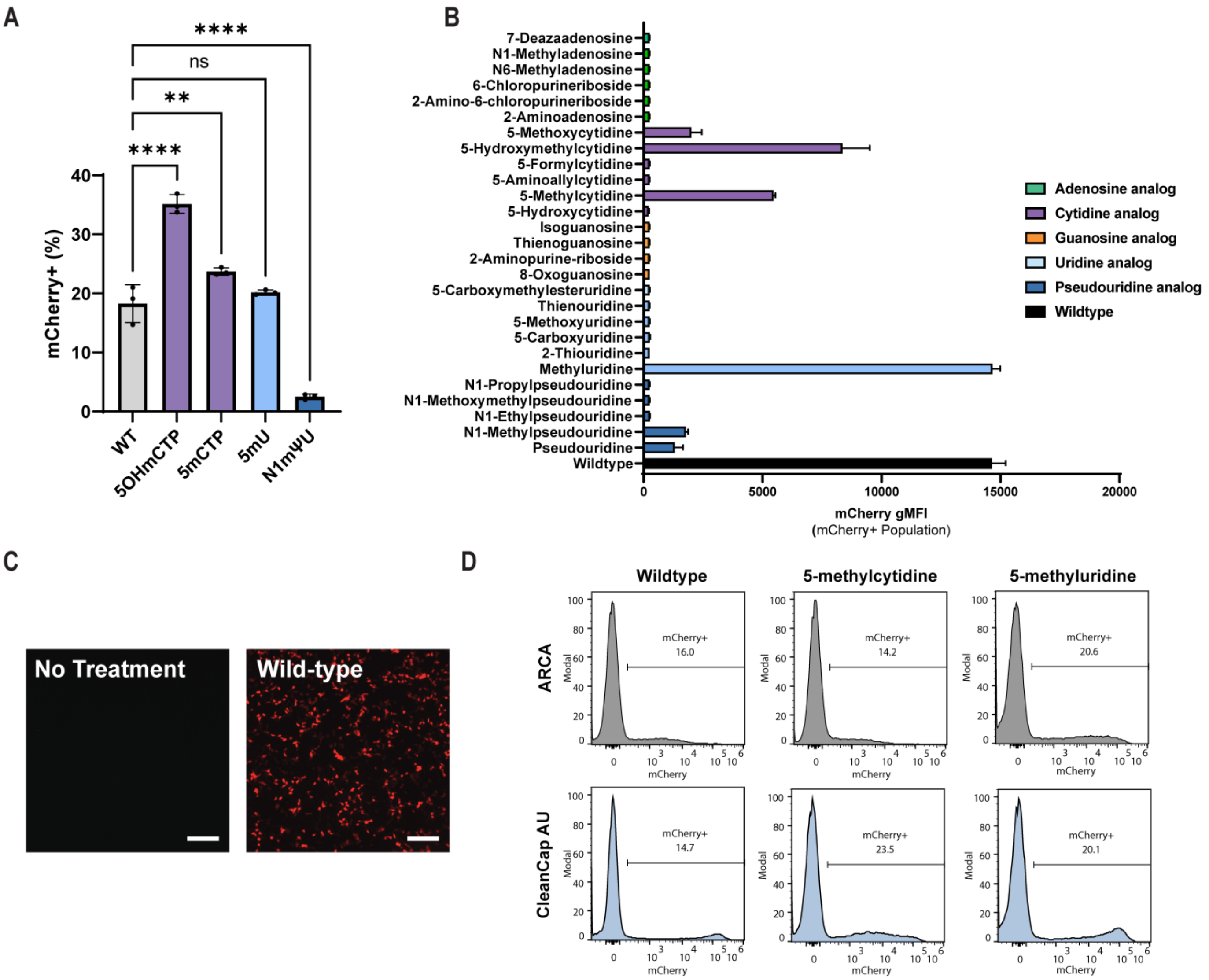


**Supplementary Figure 1.) A)** Comparison of transfection efficiency of HEK293-T cells transfected with modified saRNA relative to unmodified saRNA. Error bars indicate standard deviation from n = 3 biological replicates. **B)** Geometric mean mCherry fluorescence of HEK293-T cells transfected with the library of modified saRNA. **C)** Live-cell microscopy images of control samples from Figure 1D. Wildtype refers to unmodified saRNA. **D)** Representative flow cytometry histograms of HEK293-T cells transfected with modified saRNA synthesized with ARCA (top row) or CleanCap AU (bottom row).

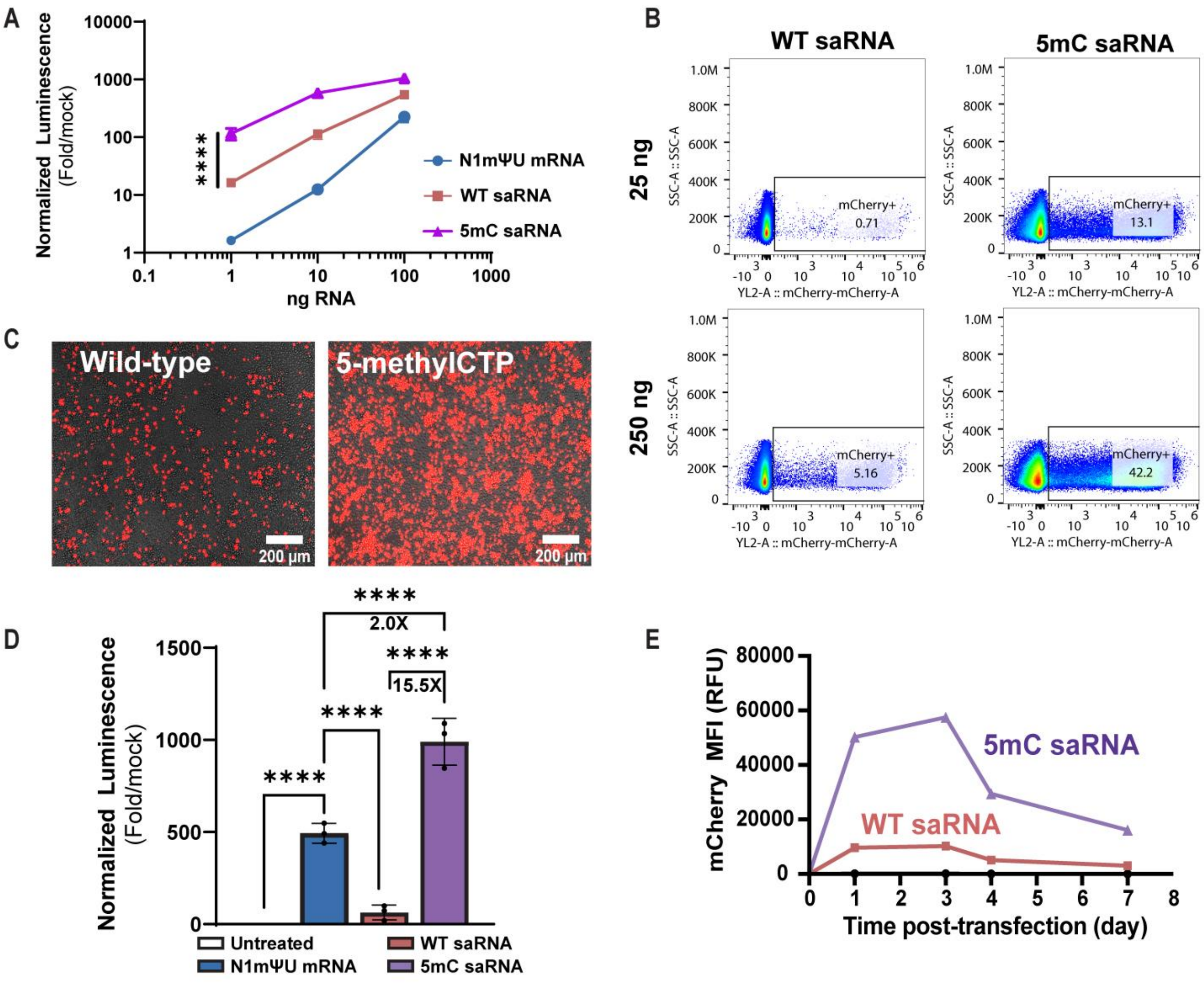


**Supplementary Figure 2.) A)** Dose response of HEK293-T transfected with LNPs containing N1mΨ mRNA, WT saRNA, or 5mC saRNA encoding luciferase. n = 4 biological replicates. **B)** Representative flow plots of Jurkat cells transfected with WT or 5mC saRNA encoding mCherry. **C)** Live-cell microscopy of Jurkat cells transfected with WT or 5mC saRNA encoding mCherry. **D)** Expression levels 24 hours after LNP transfection of 250 ng of mRNA or saRNA encoding a luciferase reporter in Jurkat cells. Luciferase signal is presented as fold-change compared to untransfected mock cells. n = 3 biological replicates. **E)** Median fluorescence intensity (MFI) of Jurkat cells transfected with WT or 5mC saRNA encoding mCherry over 7 days. n = 3 biological replicates.

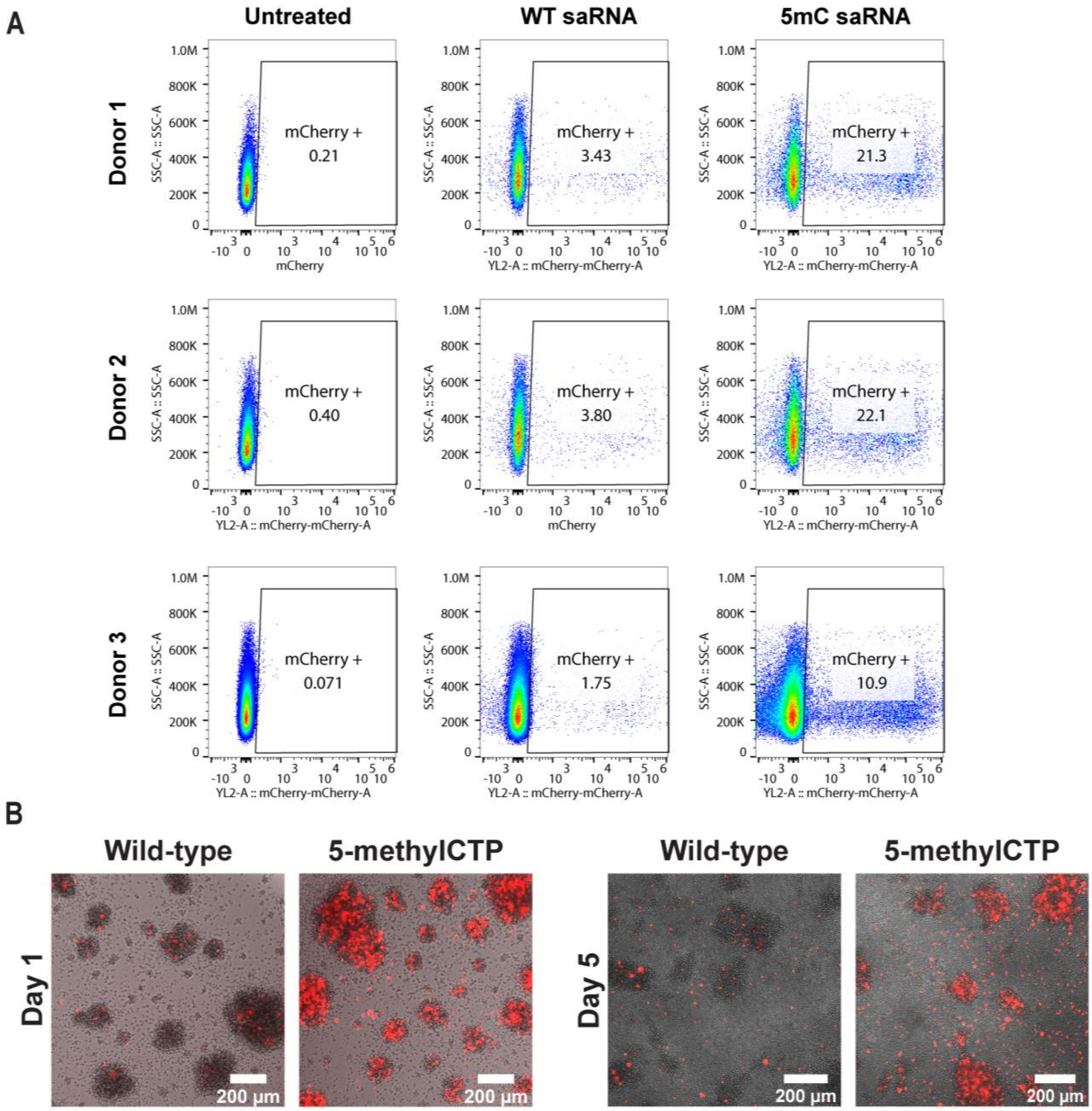


**Supplementary Figure 3.) A)** Representative flow plots of primary T cells from three different donors transfected with WT or 5mC saRNA encoding mCherry. **B)** Live cell microscopy of primary T cells transfected with WT or 5mC encoding mCherry 1 day or 5 days after transfection.

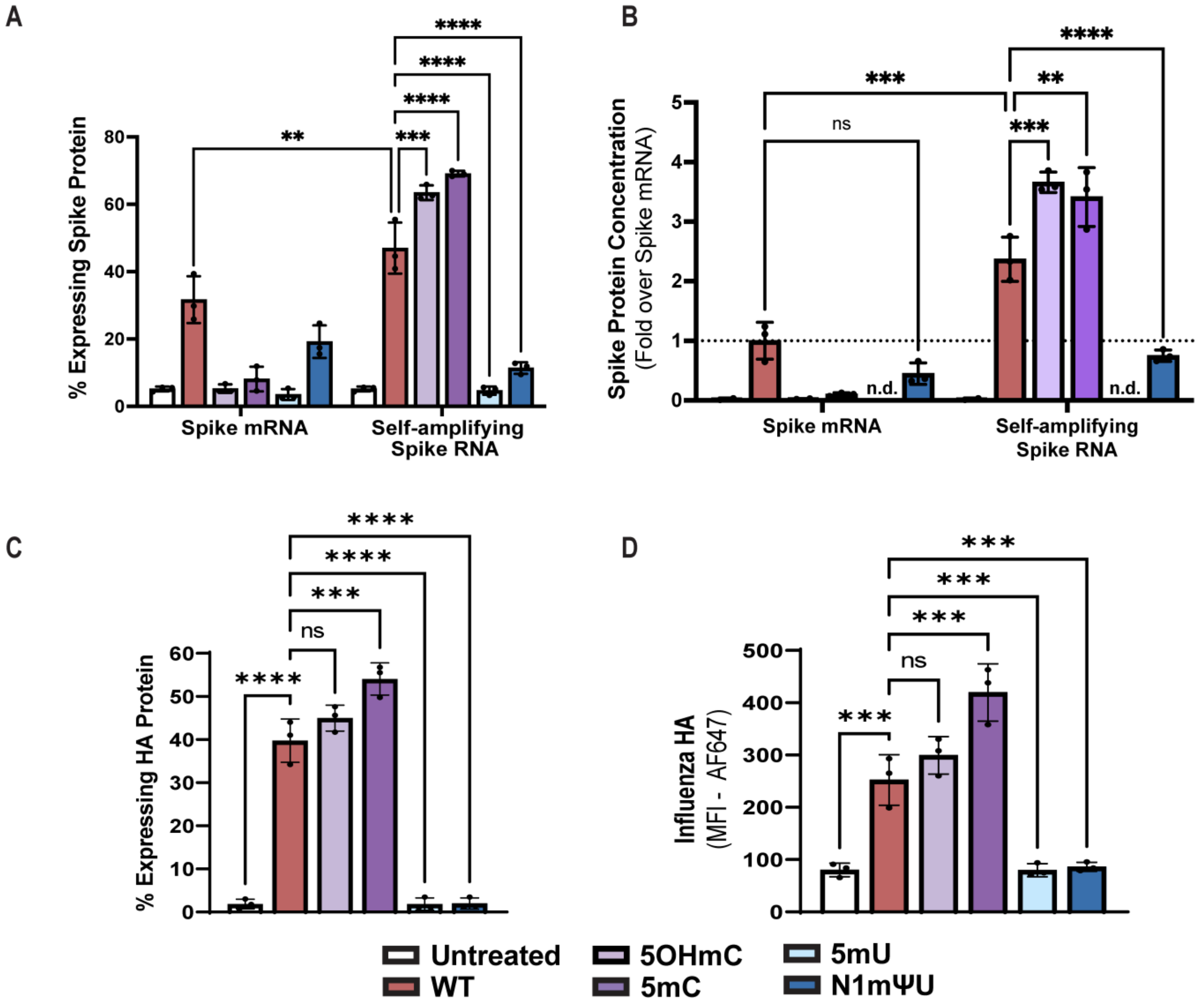


**Supplementary Figure 4.) A)** Transfection efficiency assessed by flow cytometry 24 hours after LNP transfection of C2C12 with 100 ng of modified Spike encoding mRNA or saRNA. n = 3 biological replicates. Cells were stained with an anti-Spike AF647 antibody. **B)** Detection of Spike protein by ELISA in lysed C2C12 cells after transfection with 100 ng modified Spike encoding mRNA or saRNA. Error bars represent standard deviation from n = 3 biological replicates. **C)** Transfection efficiency 24 hours after transfection of C2C12 with 25 ng of modified HA encoding mRNA or saRNA. Error bars represent standard deviation from n = 3 biological replicates. **D)** Median fluorescence intensity (MFI) of anti-HA AF647 staining in C2C12 cells. Error bars represent standard deviation from n = 3 biological replicates.

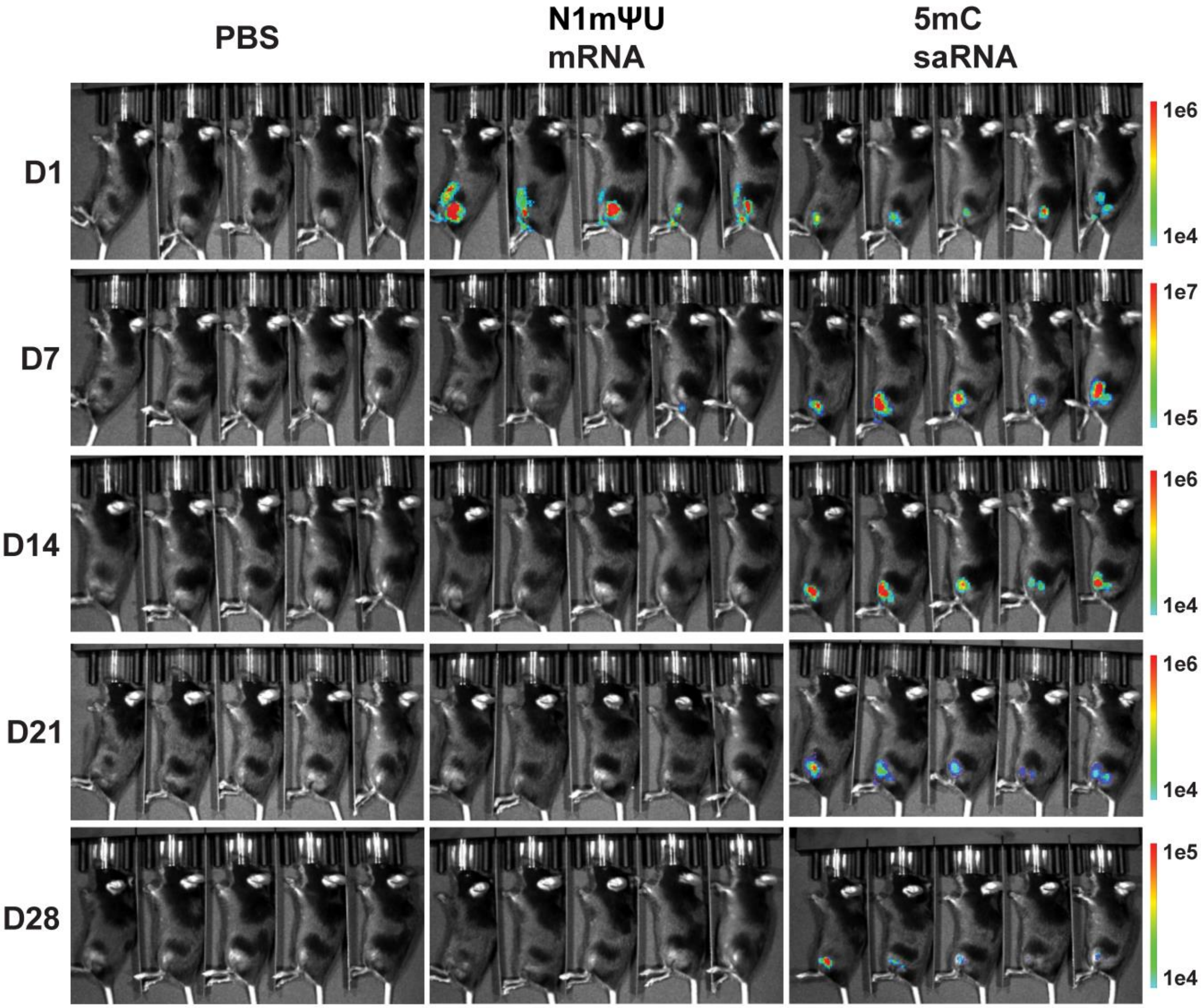


**Supplementary Figure 5.)** Whole body BLI images of mice injected in the left hind limb with 50 µL of PBS or LNPs containing 2.5 µg of mRNA or saRNA. n = 5 biological replicates.

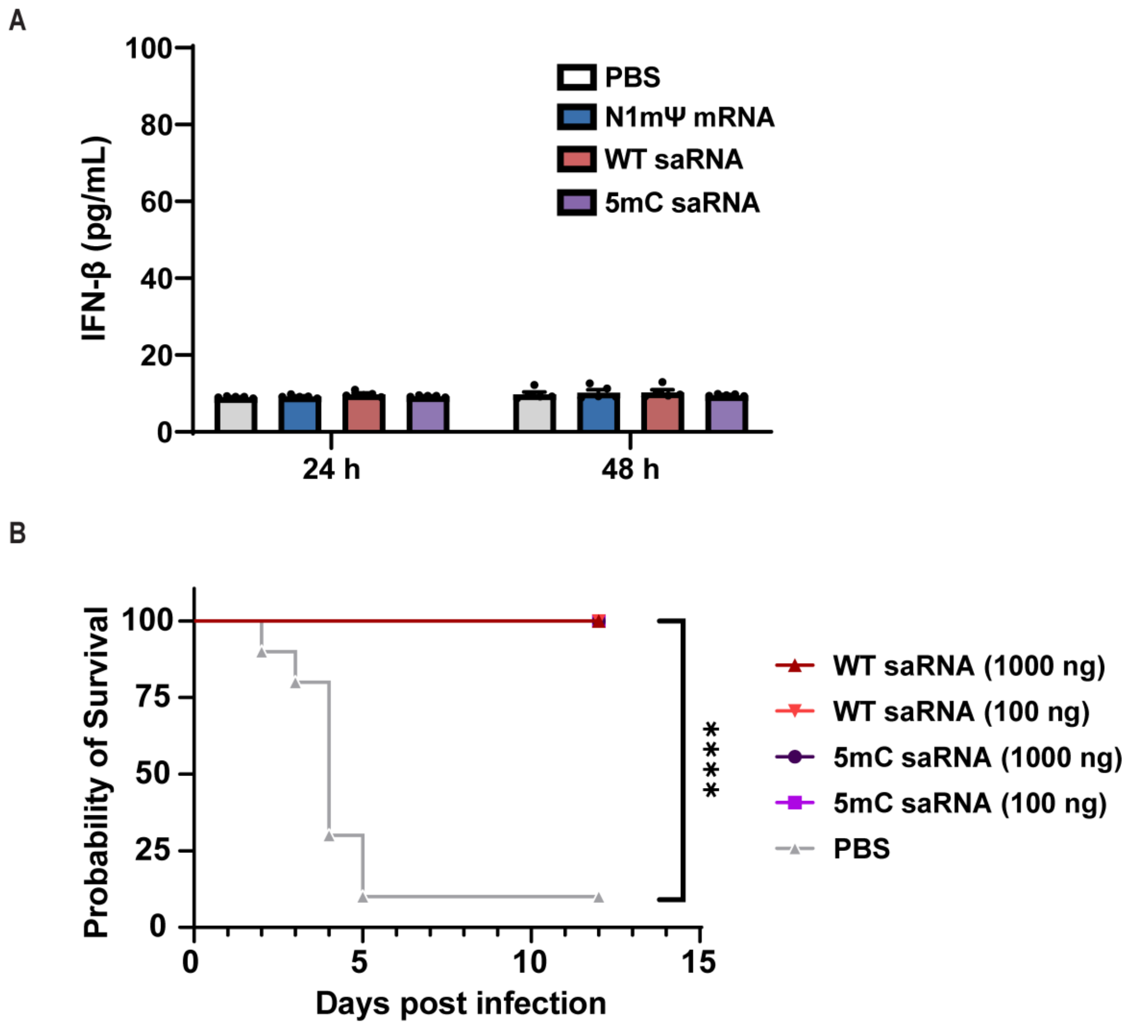


**Supplementary Figure 6.) A)** IFN-β expression in serum collected from mice 24 hours or 48 hours after initial vaccination with 1000 ng of RNA in LNPs. Error bars represent the standard error mean of n = 5 biological replicates. **B)** Survival of mice after challenge with a lethal challenge of MA30 virus. n = 10 mice per group. For survival study statistics, a log-rank (Matel-Cox) test was used between groups. **** = p < 0.0001.

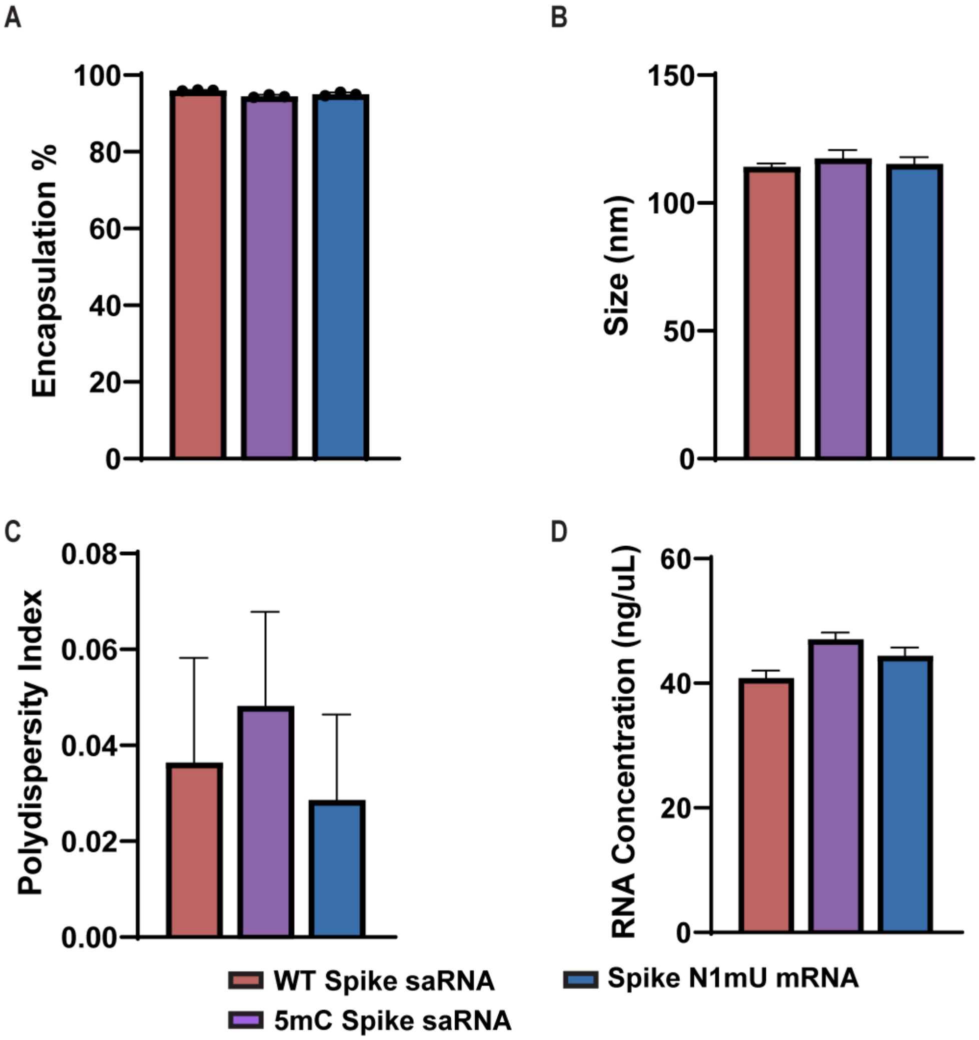


**Supplementary Figure 7.) A)** Encapsulation efficiency of post-dialysis SARS-CoV-2 spike encoding LNPs used in vaccination study. Error bars represent standard deviation from n = 3 replicate measurements per sample. **B)** Size of LNPs used in vaccination study determined by dynamic light scattering (DLS). Error bars represent standard deviation from n = 5 measurements per sample. **C)** Polydispersity index (PDI) of LNPs used in vaccination study. Error bars represent standard deviation from n = 5 measurements per sample.

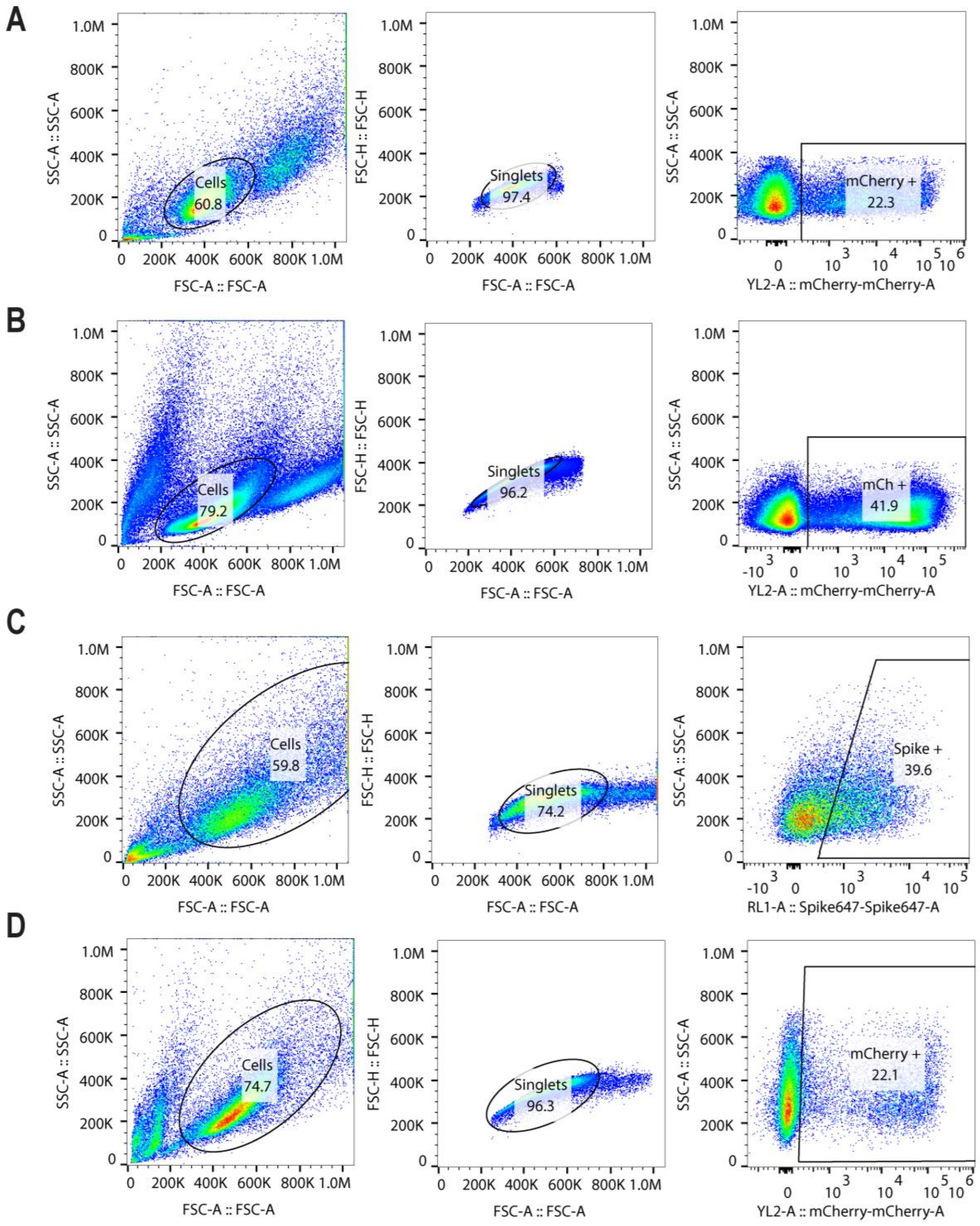


**Supplementary Figure 8.) A)** Gating strategy used in screening of modNTP saRNA library via HEK293-T transfection. **B)** Gating strategy used in analysis of transfection efficiency of saRNA LNPs in Jurkat T cells. **C)** Gating strategy for determining the expression of SARS-CoV-2 Spike protein in HEK or C2C12 cells. **D)** Gating strategy used in analysis of transfection efficiency of saRNA LNPs in primary T cells.
